# Regulation of food intake by astrocytes in the brainstem dorsal vagal complex

**DOI:** 10.1101/543991

**Authors:** Alastair J. MacDonald, Fiona E. Holmes, Craig Beall, Anthony E. Pickering, Kate L.J. Ellacott

## Abstract

Food intake is controlled by the coordinated action of numerous brain regions but a complete understanding remains elusive^1^. Of these brain regions the brainstem dorsal vagal complex (DVC) is the first site for integration of visceral synaptic and hormonal cues that act to inhibit food intake^2^. The DVC consists of three nuclei: the nucleus of the solitary tract (NTS), area postrema (AP) and dorsal motor nucleus of the vagus (DMX). Targeted chemogenetic activation of appetite-responsive NTS neuronal populations causes short term decreases in food intake^3–7^. Astrocytes are a class of glial cell which provide metabolic and structural support to neurons and play an active role in modulating neurotransmission. Within the hypothalamic arcuate nucleus (ARC) astrocytes are regulated by both positive and negative energy balance and express receptors for hormones that influence satiety and hunger^8–11^. Chemogenetic activation of these ARC astrocytes alters food intake^11–13^. Since NTS astrocytes respond to vagal stimulation^14^, we hypothesised that they may be involved in mediating satiety. Here we show that NTS astrocytes show plastic alterations in morphology following excess food consumption and that chemogenetic activation of DVC astrocytes causes a decrease in food intake, by recruiting an appetite-inhibiting circuit, without producing aversion. These findings are the first using genetically-targeted manipulation of DVC astrocytes to demonstrate their role in the brain’s regulation of food intake.

In order to examine whether NTS astrocytes respond to changes in energy status we induced short-term positive energy balance by allowing mice to exclusively eat a high-fat chow for 12 hours during the dark-phase. This paradigm induced a feeding binge when compared with standard chow-fed mice (Fig. 1a). High-fat fed mice had a greater number of glial fibrillary acidic protein (GFAP)-immunoreactive astrocytes within the NTS when compared with controls, with the changes being most pronounced in the NTS at the level of the AP (Fig. 1c-e). GFAP-immunoreactive astrocytes in the NTS adjacent to the AP of high-fat fed mice had greater morphological complexity, as assessed by Sholl analysis^15,16^, and a greater number of processes than those of standard chow fed controls (Fig. 1f,g). These findings suggest that NTS astrocytes show dynamic reactive changes to the acute nutritional excess caused by consumption of a high-fat diet.

**Fig. 1.**
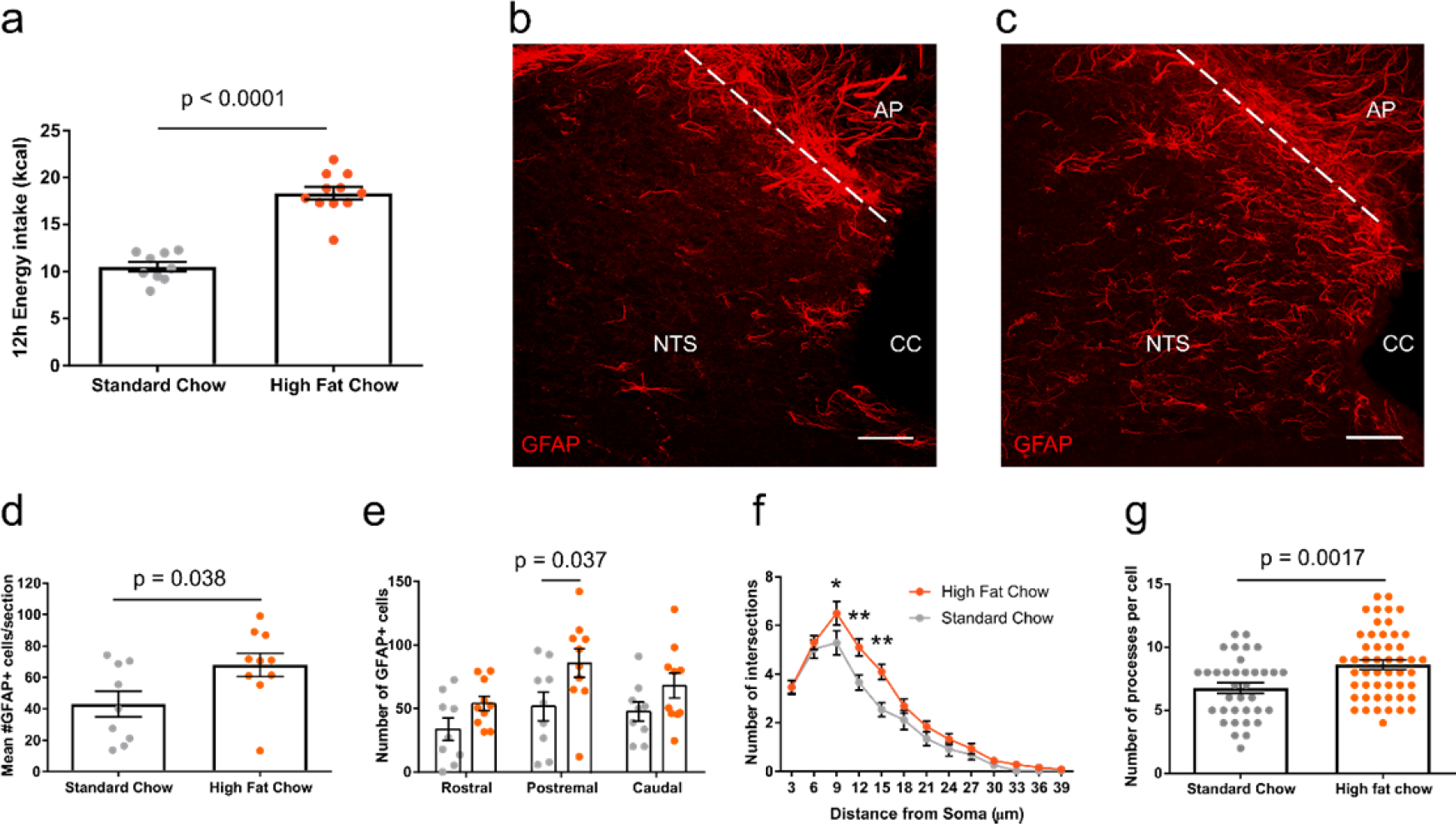
High-fat chow intake increases the number and morphological complexity of astrocytes in the nucleus of the solitary tract (NTS). **a,** Dark-phase energy intake of standard and high fat chow fed mice (10.53 ± 0.51 vs 18.37 ± 0.67 kcal, n=9-10 mice/group, p<0.0001, unpaired t-test). **b,c,** Representative maximum projection confocal image of GFAP immunostaining from a standard **(b)** and high fat **(c)** chow fed mouse, scale bar = 50 μm. **d,** Mean number of GFAP positive cells from tissue sections of NTS from standard and high fat chow fed mice (43.09 ± 8.19 vs 67.9 ± 7.47 cells, n=9-10 mice/group, p=0.038, unpaired t-test). **e,** Number of GFAP positive cells within anatomical subdivisions of NTS from standard (grey) and high-fat (orange) chow fed mice (n=9-10 mice/group, Two-way Analysis of Variance [ANOVA], Food, p=0.0018, F_(1,51)_=10.81; Rostrocaudal position, p=0.034, F_(2,51)_=3.63; interaction, p=0.7, F_(2,51)_=0.36; Sidak’s post-hoc test). **f,** Mean Sholl profile of postremal NTS astrocytes of standard and high fat chow fed mice (n=35-50 cells from 4-5 mice/group, Two-way ANOVA, Food, p<0.0001, F_(1,1079)_=23.24; Distance from soma, p<0.0001, F_(12,1079)_=108.3; interaction, p=0.04, F_(12,1079)_=1.83; Sidak’s post-hoc test). **g,** Number of processes of individual postremal NTS astrocytes from standard and high fat chow fed mice (n=35-50 cells from 4-5 mice/group, p=0.0017, unpaired t-test). * = p<0.05, ** = p<0.01. AP = area postrema, cc = central canal.

It is established that Gq-coupled designer receptors exclusively activated by designer drugs (DREADDs) can be used to selectively activate astrocytes by driving increases in intracellular Ca^2+^, the main signalling modality of these cells^11,17–19^. We bilaterally injected the DVC of mice with adeno-associated viral (AAV) vectors containing hM3Dq_mCherry (DVC::GFAP^hM3Dq^ mice) or mCherry (DVC::GFAP^mCherry^ mice) under the control of the human GFAP promoter (hGFAP) with the goal of limiting expression to DVC astrocytes (Fig. 2a,b). Examining mCherry immunoreactivity showed strong expression in the DVC (Fig. 2c). The mCherry expressing cells were immunopositive for GFAP (Figure 2d) but not the neuronal marker NeuN (Fig. 2e) indicating cell type-specificity.

**Fig. 2.**
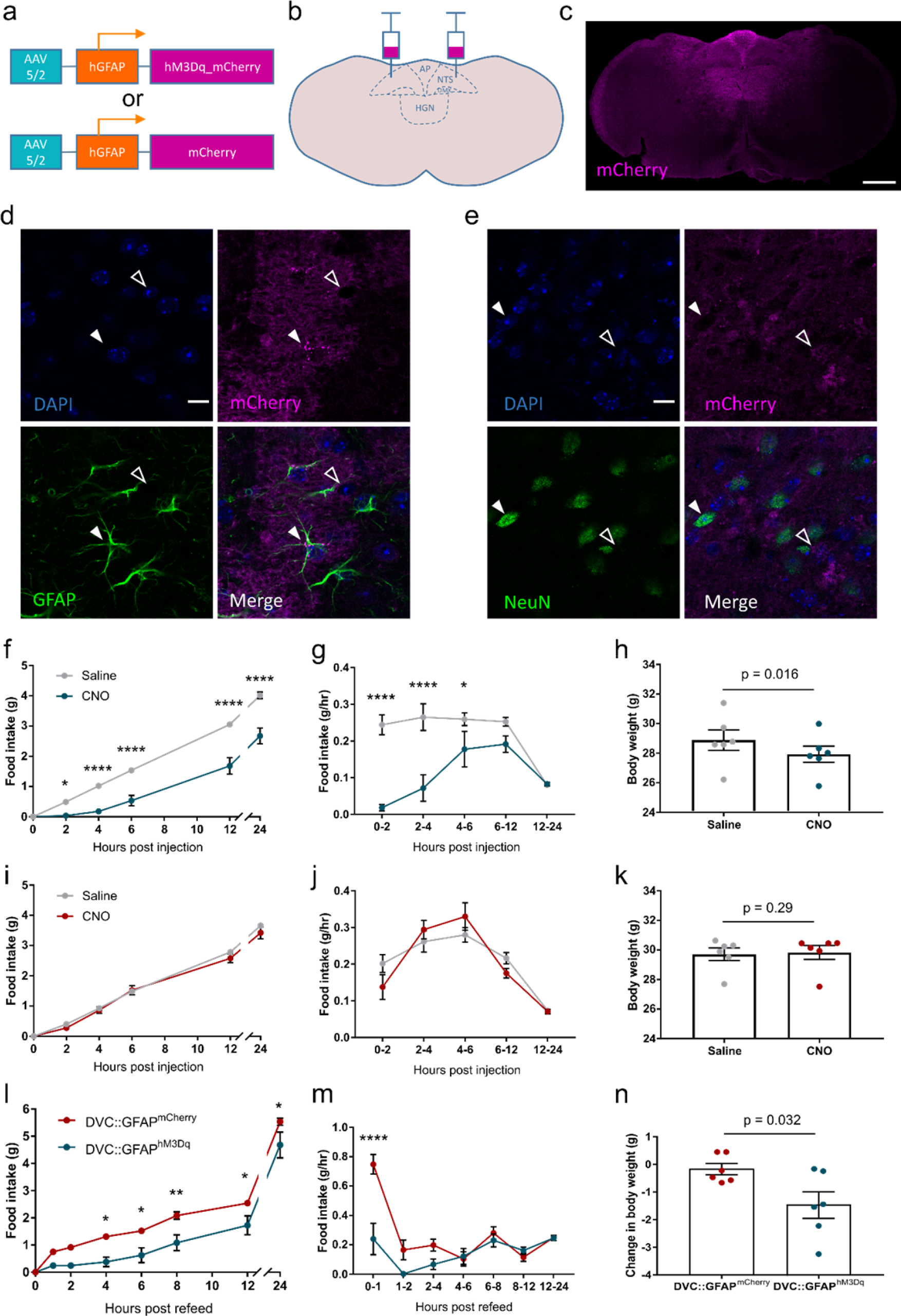
Chemogenetic activation DVC astrocytes reduces food intake. **a,** AAV vectors containing either hM3Dq_mCherry or mCherry under the hGFAP promoter. **b,** Diagram showing bilateral injection of the virus into the DVC. **c,** Representative image showing mCherry immunofluorescence in a DVC::GFAP^hM3Dq^ mouse, scale bar = 500 μm. **d,e,** Immunoreactivity for GFAP **(d)** or NeuN **(e)** and mCherry in a DVC::GFAP^hM3Dq^ mouse. Closed arrow in **(d)** shows a GFAP-positive cell, open arrow shows a GFAP-negative putative neuron while closed arrow in **(e)** shows a NeuN-positive neuron, open arrow shows a NeuN-negative putative astrocyte. Scale bar = 25 μm. **f-h,** DVC::GFAP^hM3Dq^ mice (n=6) were injected with saline or CNO 30 minutes prior to the beginning of the dark-phase. **f,** Cumulative food intake (Two-way repeated measure [RM] ANOVA, Drug, p=0.0007, F_(1,5)_=55.7; Time, p<0.0001, F_(5,25)_=331.3; interaction, p<0.0001, F_(5,25)_=15.64, Sidak’s post-hoc test). **g,** Rate of food intake (n=6 mice, Two-way RM ANOVA, Drug, p=0.0009, F_(1,5)_=49.15; Time, p=0.0001, F_(4,20)_=9.94; interaction, p<0.0001, F_(4,20)_=11.07, Sidak’s post-hoc test). **h,** Body weight 8 hours after lights-off (28.89 ± 0.69 vs 27.93 ± 0.55g, p=0.016, paired t-test). **i-k,** DVC::GFAP^mCherry^ mice (n=6) were injected with saline or CNO 30 minutes prior to the beginning of the dark-phase. **i,** cumulative food intake (Two-way RM ANOVA, Drug, p=0.33, F_(1,5)_=1.15; Time, p<0.0001, F_(5,25)_=506.7; interaction, p=0.18, F_(5,25)_=1.65, Sidak’s post-hoc test). **j,** rate of food intake (Two-way RM ANOVA, Drug, p=0.76, F_(1,5)_=0.10; Time, p<0.0001, F_(4,20)_=29.57; interaction, p=0.07, F_(4,20)_=2.56, Sidak’s post-hoc test). **k,** Body weight 8 hours after lights-off (29.59 ± 0.43 vs 29.83 ± 0.47g, p=0.29, paired t-test). **l-n,** DVC::GFAP^mCherry^ and DVC::GFAP^hM3Dq^ mice (n=6/group) were fasted for 12 hours during the dark phase then injected with CNO 30 minutes prior to reintroduction of food at the onset of the light phase. **l,** Cumulative food intake (Two-way ANOVA, Genotype, p=0.01, F_(1,10)_=10.03; Time, p<0.0001, F_(7,70)_=310.5; interaction, p=0.0053, F_(7,70)_=3.20, Sidak’s post-hoc test). **m,** Rate of food intake (Two-way ANOVA, Genotype, p=0.007, F_(1,10)_=11.4; Time, p<0.0001, F_(6,60)_=18.55; interaction, p<0.0001, F_(6,60)_=8.35, Sidak’s post-hoc test). **n,** Change in body weight from before fasting to 8 hours after refeeding (−.017 ± 0.21 vs −1.47 ± 0.48 g, p=0.024 unpaired t-test). * = p<0.05, ** = p<0.01, **** = p<0.0001. AP = area postrema, DMX = dorsal motor nucleus of the vagus, HGN = hypoglossal nucleus, NTS = nucleus of the solitary tract.

Mice typically eat most of their food during the dark-phase. Chemogenetic activation of the transduced astrocytes in DVC::GFAP^hM3Dq^ mice (clozapine-N-oxide CNO; 0.3.mg/kg, i.p immediately prior to onset of the dark-phase) produced an 83.9% reduction in food intake (4 hours after injection) compared to saline control (Fig 2f). This anorectic effect lasted approximately 6 hours after which the rate of food intake was not significantly different to the control group (Fig. 2g). Body-weight, measured 8 hours after lights-off was reduced by 4.4% on the day of CNO injection compared with the day of saline injection (Fig. 2h). In DVC::GFAP^mCherry^ mice the same CNO injection protocol had no effect on dark-phase food intake, feeding rate, or body-weight (Fig. 2i-k). Following i.p. injection of CNO, both food intake and food seeking behaviour were lower in DVC::GFAP^hM3Dq^ mice compared with DVC::GFAP^mCherry^ mice (Supplementary Fig. 1a-c). This suggests that the reduced food intake is a result of suppressed drive to feed rather than reflecting a motor impairment of feeding, such as may be anticipated as a result of activation of astrocytes in the hypoglossal nucleus (HGN) disrupting tongue movement.

To evaluate whether chemogenetic activation of DVC astrocytes was still sufficient to impact feeding when there was an increased drive to eat, we utilised a fast-induced re-feeding paradigm. CNO injection (0.3.mg/kg i.p.) prior to reintroduction of food after a 12 hour fast lowered cumulative food intake in DVC::GFAP^hM3Dq^ mice compared with DVC::GFAP^mCherry^ controls (Fig. 2l). During the first hour of refeeding, the rate of food intake was 62.7% lower in DVC::GFAP^hM3Dq^ mice, compared with DVC::GFAP^mCherry^ controls (measured after 1 hour), indicating that activation of DVC astrocytes was sufficient to acutely supress the drive to eat induced by fasting (Fig. 2m). No compensatory/rebound hyperphagia was observed in DVC::GFAP^hM3Dq^ mice in the 24 hours following CNO injection (Fig. 2l,m). In line with the differences in food intake, DVC::GFAP^mCherry^ mice recovered more of their body weight lost as a result of the fast within 8 hours of food being reintroduced than DVC::GFAP^hM3Dq^ mice (Fig. 2n).

Reductions in food intake in mice can be indicative of malaise and/or aversion^20^. To test whether DVC astrocyte activation is aversive we used a conditioned place assay (Fig 3a). During initial assessment, prior to conditioning, neither DVC::GFAP^hM3Dq^ nor DVC::GFAP^mCherry^ mice showed a preference for either chamber (Fig 3c,d). Consistent with an absence of aversion/negative salience associated with malaise, following conditioning (pairing each side of the apparatus with either saline or i.p. CNO injection) mice still showed no preference for or avoidance of a particular side of the apparatus (Fig. 3b-d). Furthermore, during conditioning trials there was no statistically significant difference in total distance travelled following saline or CNO treatment in the DVC::GFAP^hM3Dq^ or DVC::GFAP^mCherry^ groups suggesting that CNO-mediated DREADD activation did not cause malaise, sedation or impact locomotion (Fig. 3e,f).

**Fig. 3.**
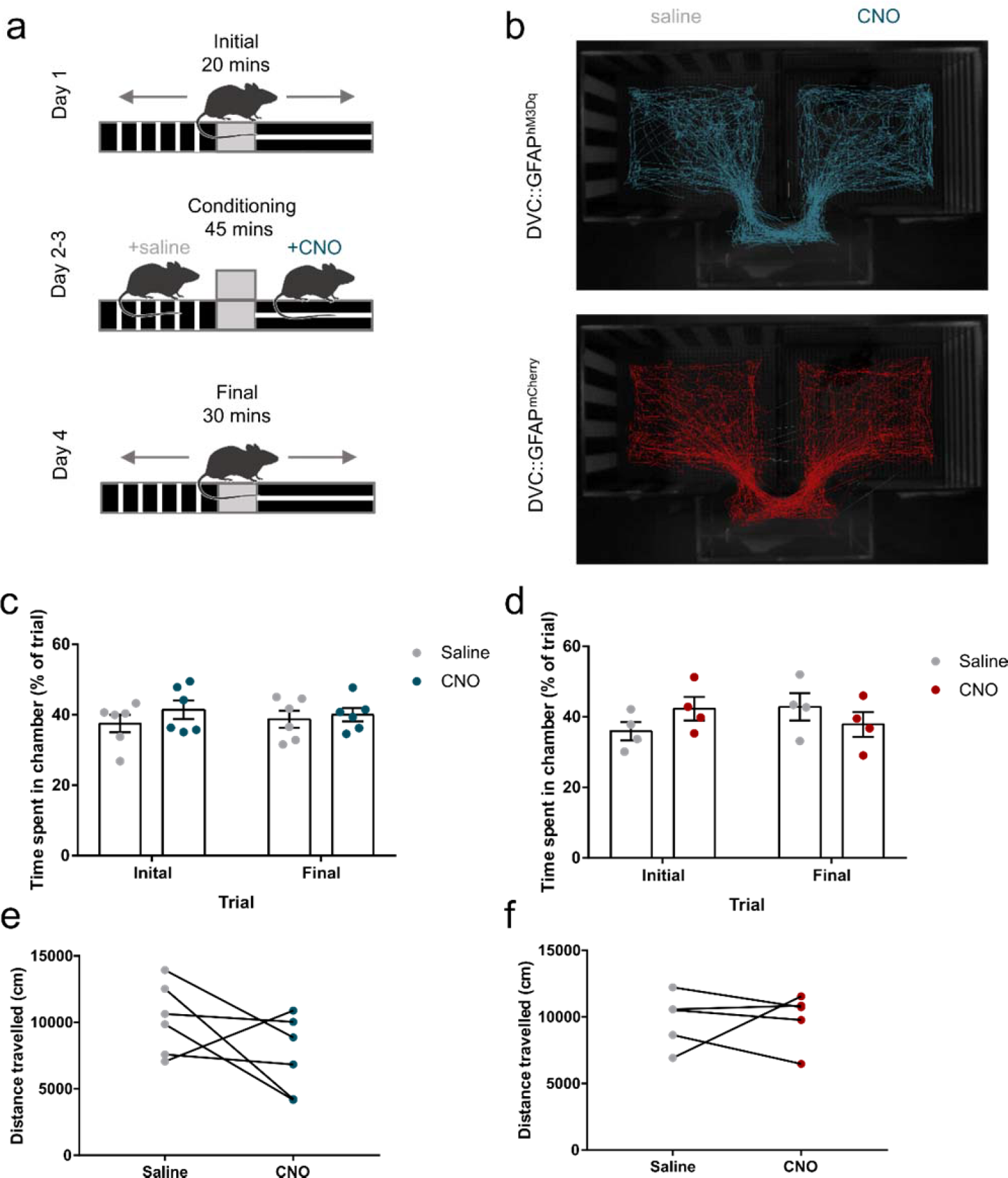
Injection with CNO does not induce conditioned place preference/aversion or affect locomotion in DVC::GFAP^hM3Dq^ or DVC::GFAP^mCherry^ mice. **a,** Schematic of the conditioning protocol. **b,** Representative tracks of a DVC::GFAP^hM3Dq^ (top, blue) and a DVC::GFAP^mCherry^ (bottom, red) mouse during the final trial. **c,** Percentage of the trial spent in the saline- and CNO-paired chamber before (initial) and after (final) conditioning sessions in DVC::GFAP^hM3Dq^ mice (n=6 mice, Two-way RM ANOVA, Drug, p=0.32, F_(1,10)_=1.09; Conditioning, p=0.96, F_(1,10)_=0.002; interaction, p=0.57, F_(1,10)_=0.35). **d,** Percentage of the trial spent in the saline- and CNO-paired chamber before and after conditioning sessions in DVC::GFAP^mCherry^ mice. (n=4 mice, Two-way RM ANOVA, Drug, p=0.76, F_(1,6)_=0.10; Conditioning, p=0.78, F_(1,6)_=0.08; interaction, p=0.23, F_(1,6)_=1.81) **e,** Distance travelled during conditioning sessions for DVC::GFAP^hM3Dq^ mice (10260 ± 1101 vs 7501±1183 cm, n=6 mice, p=0.18, paired t-test). **f,** Distance travelled during conditioning sessions for DVC::GFAP^mCherry^ mice (9773±991.1 vs CNO 9870±896.3 cm, n=5 mice, p=0.94, paired t-test).

To confirm DVC circuit activation and to examine putative downstream targets, mice were injected with CNO (0.3.mg/kg i.p.) 2-3 hours prior to perfusion and immunohistochemistry for the immediate early gene c-FOS, as a marker of neuronal activation. The number of c-FOS expressing cells was greater in DVC::GFAP^hM3Dq^ mice compared with DVC::GFAP^mCherry^ control mice in the NTS and AP (Fig. 4a-e), and the downstream target of NTS neurons the lateral parabrachial nucleus (lPBN) (Fig. 4f-i). However in the paraventricular nucleus of the hypothalamus (PVH), another projection target of satiety signalling NTS neurons^5,21^, there was no difference in the number of c-FOS expressing cells between groups (Fig. 4j-m). This indicates that CNO activates astrocytes in DVC::GFAP^hM3Dq^ mice which leads to the recruitment of local and downstream neuronal circuitry implicated in regulating energy homeostasis^22,23^.

**Fig. 4.**
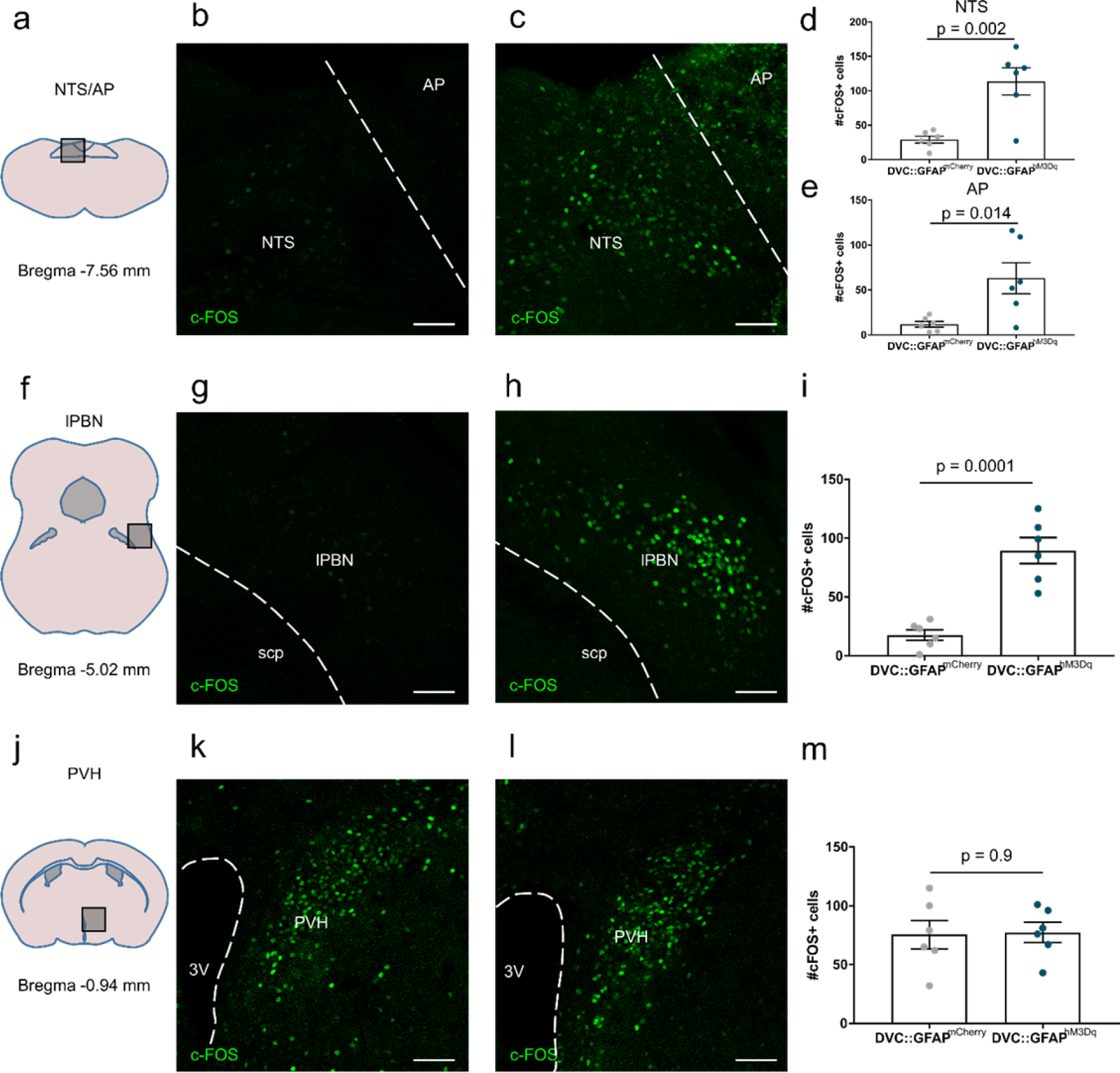
Injection with CNO induces c-FOS-immunoreactivity in a downstream neural target in DVC::GFAP^hM3Dq^ mice. DVC::GFAP^mCherry^ and DVC::GFAP^hM3Dq^ mice (n=6 mice/group) were injected with CNO 2-3 hours prior to perfusion. **a-c,** Representative images of c-FOS immunostaining from the DVC of a DVC::GFAP^mCherry^ mouse **(b)** and a DVC::GFAP^hM3Dq^ mouse **(c). d,e,** Quantification of c-FOS immunoreactive cells in NTS **(d)** (29 ± 5.01 vs 113.7 ± 19.62 cells, p=0.002, unpaired t-test) and AP **(e)** (11.83 ± 3.2 vs 63.17 ± 17.2 cells, p=0.014, unpaired t-test). **f-h,** Representative images of c-FOS immunostaining from the lPBN of a DVC::GFAP^mCherry^ mouse **(g)** and a DVC::GFAP^hM3Dq^ mouse **(h). i,** Quantification of c-FOS immunoreactive cells in lPBN (17.5 ± 4.49 vs 89.33 ± 11.08 cells, p=0.002, unpaired t-test). **j-l,** Representative images of c-FOS immunostaining from the lPBN of a DVC::GFAP^mCherry^ mouse **(k)** and a DVC::GFAP^hM3Dq^ mouse **(l). m,** Quantification of c-FOS immunoreactive cells in PVH (75.5 ± 12.05 vs 77.33 ± 8.58 cells, p=0.90, unpaired t-test). For all images scale bar = 50 μm. AP = area postrema, NTS = nucleus of the solitary tract, lPBN = lateral parabrachial nucleus, scp = superior cerebellar peduncle, PVH = paraventricular nucleus of the hypothalamus.

In this study we have shown that astrocytes in the NTS react to excess intake of an energy dense food by upregulating GFAP expression and show morphological plasticity. Furthermore, DREADD-mediated activation of astrocytes in the DVC causes a potent but reversible decrease in food intake, associated with activation of local and distant neuronal circuits, without inducing emotionally salient associations. This evidence supports the hypothesis that these astrocytes are involved in a homeostatic response to increases in food intake and their activation acts to drive a counter-acting decrease in food intake to restore energy balance.

Astrocytes of the hypothalamic arcuate nucleus (ARC) have also been implicated in the regulation of food intake^8^. In the hypothalamus these glia are activated by short-term energy imbalance and direct manipulation of their activity using DREADDs alters food intake^8,11,12,24^. The first study using DREADDs to activate ARC astrocytes from Yang and colleagues observed a decrease in food intake during the dark-phase of a smaller magnitude and shorter time duration than we observed with DVC-astrocyte activation^12^. However a second group, Chen and colleagues, observed the converse effect: an increase in dark-phase feeding with a short (~1hr post injection) time course^11^. Both studies reported that CNO injection drove feeding behaviour in the light-phase but again Chen et al showed an immediate (within 30 min of injection) effect while Yang et al showed a delayed effect (increased cumulative intake at 9 hrs after injection). It appears that chemogenetic DVC astrocyte activation has effects on feeding distinct from those described following comparable studies examining ARC astrocyte activation with respect to the magnitude, direction and onset/duration of the effect.

Our study builds upon evidence showing astrocytes of the NTS sense hormonal satiety signals^25^ and are involved in integration of vagal neurotransmission^14^, but importantly extends these studies by using direct, inducible manipulation of these cells and studying behaviour in freely moving animals.

What remains unclear is the mechanism(s) by which astrocytes are activated in the state of caloric excess. Likely candidates include vagal-derived glutamate^14^, Glucagon-like peptide-1 (GLP-1) mediated signalling^25^ and circulating fatty-acids driving local inflammation. These factors may act in combination to drive activation of astrocytes in the NTS. NTS astrocytes directly sense vagal glutamate release via Ca^2+^-permeable AMPA receptors expressed on the cell membrane^14^. Furthermore, in rats, NTS astrocytes take up exendin-4, an agonist of the GLP-1R. Metabolic inhibition of astrocytes neighbouring the 4V abolishes the anorexigenic effect of exendin-4, suggesting these cells are responsible for mediating the effect^25^. Two studies that manipulate inflammatory nuclear factor kappa b (NF-κB) signalling in all GFAP-expressing astrocytes report feeding phenotypes, namely increased initial intake of a high-fat diet following initial exposure and resistance to the obesity phenotype when already on a high-fat diet^8,26^. Although these studies attribute the observed effects to astrocytes of the hypothalamus the contribution of NTS astrocytes, identified here as responsive to high-fat diet intake, cannot be ruled out.

The induction of c-FOS-immunoreactivity in a distal target, namely the lPBN, following CNO injection in DVC::GFAP^hM3Dq^ mice strongly suggests the recruitment of long range neuronal circuitry following DVC astrocyte activation. Since there are a number of distinct neuronal populations in the NTS whose activation induces a similar reduction in feeding, namely proopiomelanocortin^3,4^, cholecystokinin^5,6^, tyrosine hydroxylase^6^ and pre-proglucagon^7^, it is likely that these are among the neurons recruited by chemogenetic DVC astrocyte activation. Astrocytes can modulate the activity of neurons by mechanisms including altered glutamate transport and release of neuroactive molecules (e.g. ATP, d-serine). In the NTS, synaptic clearing of glutamate by astrocytes has been shown to restrain NTS neuronal firing and vagal outflow to cardiorespiratory organs^27^. As such, astrocyte glutamate transport can manipulate the firing rates of NTS neurons and alter output from the NTS. An additional mechanism of communication is active gliotransmission. Activation of a GPCR expressed on NTS astrocytes (protease activated receptors [PAR]) leads to activation of neurons by glutamate, possibly exocytosed by astrocytes^28^. Given that antagonism of NMDA receptors in the NTS increases meal size^29^ it is possible that activation of these receptors by astrocyte-derived D-serine would reduce food intake.

Our experiments show that astrocytes are a previously overlooked component of physiological DVC-mediated satiety and provide the first causal link between their activity and the regulation of feeding. We propose that NTS/DVC astrocytes are a key component in satiety signalling that act to modulate the integration of vagal and hormonal inputs to the brainstem and as such represent a potential target for intervention in obesity.

## Acknowledgements

The authors of this manuscript have no conflicts of interest to declare. This study was funded by grants from the Medical Research Council (MR/N012763/1 KLJE and CB; MR/P025749/1 to AEP), and Diabetes UK (RD Lawrence Fellowship to CB; 13/0004647), and institutional funds from the University of Exeter Medical School (CB and KLJE). AJM is a GW4 BioMed doctoral training programme student funded by Medical Research Council.

## Materials and Methods

### Mice

All animal studies were conducted in accordance with the UK Animals in Scientific Procedures Act 1986 (ASPA) and study plans were approved by the institutional Animal Welfare and Ethical Review Body at the University of Bristol and/or Exeter. Male C57BL6/J mice (Charles River) were used for all experiments. Unless stated otherwise, mice were group housed on a 12:12 light-dark cycle at 22 ± 2 °C, with unlimited access to standard laboratory rodent diet (EURodent diet [5LF2], LabDiet, UK) and water.

### Dark-phase high-fat feeding studies and histology

Two independent cohorts of mice (aged 16 weeks) were individually housed for 4-5 days. At lights-off, standard chow was substituted for high-fat chow (D12492, TestDiet, USA) with control animals maintained on standard chow (EURodent diet). 12 - 14 hours later mice were transcardially perfused under deep sodium pentobarbital anaesthesia with heparinised 0.9% saline followed by 4% paraformaldehyde. Brains were post fixed in 4% paraformaldehyde for 4-6 hours before being transferred to 30% sucrose in phosphate buffered saline (PBS). 30 μm coronal sections of the hindbrain were taken with a freezing microtome (Bright instruments, UK) in 4 series. One series was taken and washed 3 times in 0.01M PBS then incubated for 1 hr at room temperature in 0.01M PBS with 5% normal donkey serum (Sigma, UK) and 0.3% triton X-100 (T8787, Sigma, UK). Following this, sections were incubated overnight at 4°C in mouse anti-GFAP (MAB360, Millipore, UK) diluted 1:5000 in 0.01M PBS with 1% normal donkey serum and 0.3% triton X-100. Sections were washed 8 times in 0.01M PBS and incubated in for 1 hr at room temperature donkey anti-mouse Alexa Fluor 568 (A10037, Invitrogen, UK) diluted 1:500 in 0.01M PBS with 0.3% triton X-100. Sections were washed 8 times in 0.01M PBS, mounted onto glass slides and coverslipped with fluoroshield mounting medium with DAPI (ab104139, Abcam, UK). Images for counting were acquired at 10X magnification on a fluorescence microscope (DM 4000, Leica, Germany). NTS subdivisions (Fig. 1e) were calculated as follows: rostral = −7.08mm −7.2mm from Bregma, postremal = −7.32mm − 7.64mm from Bregma, cadual = −7.76mm − 8mm from Bregma. Images for morphological analysis were acquired at 20X on a confocal microscope (DMi8, Leica, Germany). Images were analysed in FIJI^30^ (NIH, USA) using the cell counter, simple neurite tracer and Sholl analysis plugins^16^. For tracing, 5 cells per image from 2 images per animal were randomly selected for tracing and morphological analysis. The investigator was blinded to the diet of the mice during staining, acquisition and analysis.

### NTS viral vector injection

Mice (8-12 weeks) were injected with adeno-associated viral vectors in the NTS as described previously^4^. The vectors used were AAV5/2-hGFAP-hM3D(Gq)_mCherry (v97-5, titre ≥ 7.5×10^12^ viral genomes/ml; University of Zurich Viral Vector Facility, Switzerland) and AAV5/2-hGFAP-mCherry (custom preparation, titre 3.89×10^13^ viral genomes/ml; ViGene Biosciences, USA) and were diluted 1:1 in sterile PBS prior to injection. In brief, mice were deeply anaesthetised with ketamine (70 mg/kg) and medetomidine (0.5 mg/kg) and placed in a stereotaxic frame (David Kopf Instruments, USA) with the nose angled down by 20°. An incision from the crest of the skull to the nape of the neck was made with a scalpel, the muscles were parted and the atlanato-occipital membrane removed to expose the surface of the brainstem. Injections were from by a pulled glass pipette attached to an injection system (Neurostar, Germany) mounted on the stereotaxic frame at an angle of 35° to the vertical, tip facing rostral. The pipette was inserted 400 μm lateral to the midline at the level of calamus scriptorius to a depth of 1mm. Four injections of 180 nl were made at depths of −1 mm, −750 μm, −500 μm and −250 μm respectively at a rate of 100 nl/minute. The pipette was left in place for 1 minute after each injection. These injections were then repeated on the contralateral side. Mice were injected with atipamezole (1 mg/kg) and buprenorphine (0.1 mg/kg) for anaesthetic reversal and analgesia respectively and transferred to a heated cage to recover. Following surgery mice were individually housed for the duration of the experiment. The investigator was blinded to mouse group allocations for all of the subsequent experiments.

### Measurement of feeding

At 4-8 weeks following surgery, mice were acclimatised to experimenter handling and intraperitoneal injections of saline for 4 days. Mice were given an intraperitoneal (i.p.) injection of saline 15-30 minutes prior to the beginning of the dark-phase. Food was removed at the time of injection and returned to cages at the onset of the dark-phase. Food intake was measured at 2, 4, 6, 12 and 24 hours after lights-off and body weight was measured 8 hours after lights-off. The following day mice were given an i.p. injection of CNO (0.3 mg/kg, 4936, Tocris, UK) and food intake and body weight measured at the same time points. For the fast-refeed experiment food was removed from cages at the beginning of the dark-phase and 15-30 minutes prior to the light phase animals were injected with 0.3 mg/kg CNO i.p. Food was returned to cages at the onset of the light phase and food intake was measured 1, 2, 4, 6, 8, 12 and 24 hours after lights-on. Body weight was measured prior to food being removed and at 8 hours after the reintroduction of food.

### Conditioned place preference assay

For conditioned place preference testing an apparatus consisting of 2 chambers joined by a clear plastic external corridor (see figure 3b) was used. The left chamber had horizontal black and white stripes on the walls and a perforate floor and the right chamber had vertical black and white stripes on the walls and a floor with horizontal grating. On the first day mice were given 20 minutes of free access to the whole apparatus. This session was recorded using a video camera and used to determine initial preferences. On the second day mice were assigned to receive an i.p. injection of either 0.3 mg/kg CNO or an equivalent volume of saline 15 minutes prior to being placed in either the left or right chamber for 45 minutes, with the access to the second chamber blocked. On the third day mice received the opposite treatment with their access restricted to the alternate chamber, compared to the second day. On the fourth and final day mice were given free access to the whole apparatus for 30 minutes. Again this session was recorded and used to determine conditioned preference. Recorded sessions were analysed offline using Ethovision XT (Noldus, Netherlands) tracking software.

### Evaluation of DREADD-induced c-FOS activation and histological analysis

Mice were injected i.p. with 0.3 mg/kg CNO and food removed from the cage. 2-3 hours later mice were transcardially perfused as described above. 30 μm coronal sections were taken and stained as described above with primary antibodies against GFAP, mCherry, NeuN and c-FOS (Table 1) and appropriate secondary antibodies donkey anti-mouse Alexa Fluor488, donkey anti-goat Alexa Fluor594 and donkey anti-rabbit Alexa Fluor488 (Invitrogen, UK). For double immunohistochemistry mCherry staining was performed first followed by GFAP or NeuN. Images were acquired on a confocal microscope (DMi8, Leica, Germany). For c-FOS quantification, images were analysed in FIJI with the cell counter plugin.

**Table 1.** Primary antibodies used

| Antibody | Dilution | Manufacturer |
| --- | --- | --- |
| Mouse anti-GFAP | 1:5000 | Merck MAB360, UK |
| Rabbit anti-NeuN | 1:2000 | AbCam ab177487, UK |
| Goat anti-mCherry | 1:1000 | Sicgen AB0040-200, Portugal |
| Rabbit anti-c-FOS | 1:2000 | Cell Signalling Technologies #2250S, UK |

### Statistical analysis

Values for all experiments were imported in to Prism 7 (Graph Pad, USA) and the appropriate statistical tests conducted. Graphs were generated in Prism and figures prepared in Inkscape (inkscape.org).

---

**Supplemental Figure 1.**
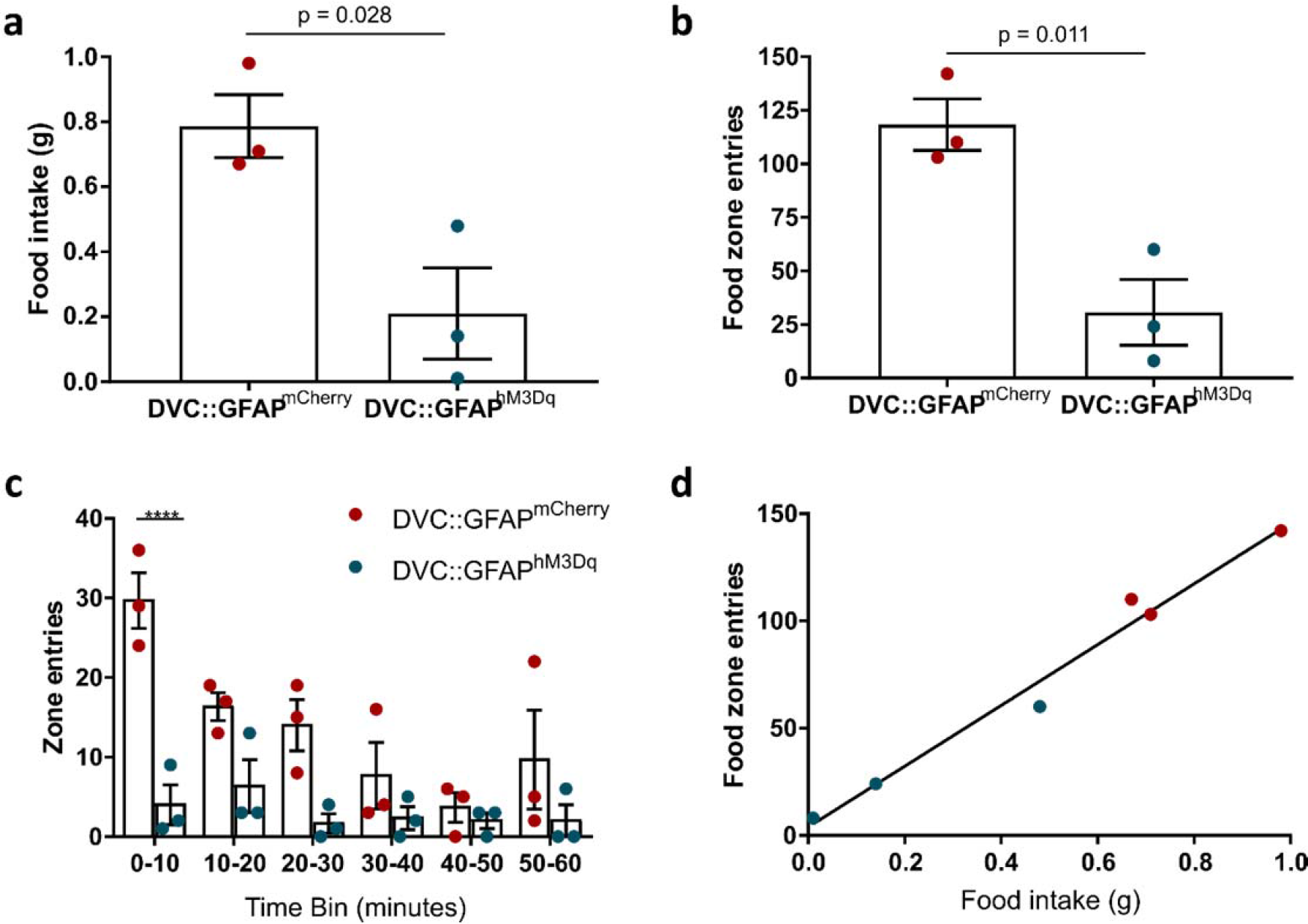
Food seeking is lower in DVC::GFAP^hM3Dq^ than DVC::GFAP^mCherry^ mice following CNO injection. DVC::GFAP^mCherry^ and DVC::GFAP^hM3Dq^ mice (n=3 mice/group) were injected with CNO at the beginning of the dark-phase and video monitored for 3 hours in their home cage with food pellets in the far corner from their nest. **a,** Food intake during the 3 hour monitoring period (p=0.028, unpaired t-test). **b,** Total number of entries to the food containing zone of the cage made during the 3 hour monitoring period (n=3 mice per group, p=0.011, unpaired t-test). **c,** Number of entries to the food containing zone of the cage in ten minute bins made by DVC::GFAP^mCherry^ (red) and DVC::GFAP^hM3Dq^ (blue) mice in the first hour of the monitoring period (Two-way ANOVA, Genotype, p=0.0052, F_(1,4)_=30.81; Time, p<0.0019, F_(5,20)_=5.74; interaction, p<0.013, F_(5,20)_=3.85, Sidak’s post-hoc test). **d,** relationship between total food intake and total entries to the food containing zone (linear regression slope = 141.8, r^2^ = 0.98). **** = p<0.0001.

## References

1. Andermann, M. L. & Lowell, B. B. Toward a Wiring Diagram Understanding of Appetite Control. Neuron 95, 757–778 (2017).

2. Grill, H. J. & Hayes, M. R. Hindbrain neurons as an essential hub in the neuroanatomically distributed control of energy balance. Cell Metab. 16, 296–309 (2012).

3. Zhan, C. et al. Acute and long-term suppression of feeding behavior by POMC neurons in the brainstem and hypothalamus, respectively. J. Neurosci. 33, 3624–3632 (2013).

4. Cerritelli, S., Hirschberg, S., Hill, R., Balthasar, N. & Pickering, A. E. Activation of brainstem proopiomelanocortin neurons produces opioidergic analgesia, bradycardia and bradypnoea. PLoS One 11, 1–26 (2016).

5. D’Agostino, G. et al. Appetite controlled by a cholecystokinin nucleus of the solitary tract to hypothalamus neurocircuit. Elife 5, 1–15 (2016).

6. Roman, C. W., Derkach, V. A. & Palmiter, R. D. Genetically and functionally defined NTS to PBN brain circuits mediating anorexia. Nat. Commun. 7, 11905 (2016).

7. Gaykema, R. P. et al. Activation of murine pre-proglucagon-producing neurons reduces food intake and body weight. J. Clin. Invest. 127, 1031–1045 (2017).

8. Buckman, L. B. et al. Evidence for a novel functional role of astrocytes in the acute homeostatic response to high-fat diet intake in mice. Mol. Metab. 4, 58–63 (2015).

9. Kim, J. G. et al. Leptin signaling in astrocytes regulates hypothalamic neuronal circuits and feeding. Nat. Neurosci. 17, 908–10 (2014).

10. Fuente-Martín, E. et al. Ghrelin Regulates Glucose and Glutamate Transporters in Hypothalamic Astrocytes. Sci. Rep. 6, 23673 (2016).

11. Chen, N. et al. Direct modulation of GFAP-expressing glia in the arcuate nucleus bi-directionally regulates feeding. Elife 5, 1–21 (2016).

12. Yang, L., Qi, Y. & Yang, Y. Astrocytes Control Food Intake by Inhibiting AGRP Neuron Activity via Adenosine A1 Receptors. Cell Rep. 11, 798–807 (2015).

13. Sweeney, P., Qi, Y., Xu, Z. & Yang, Y. Activation of hypothalamic astrocytes suppresses feeding without altering emotional states. Glia 64, 2263–2273 (2016).

14. McDougal, D. H., Hermann, G. E. & Rogers, R. C. Vagal Afferent Stimulation Activates Astrocytes in the Nucleus of the Solitary Tract Via AMPA Receptors: Evidence of an Atypical Neural-Glial Interaction in the Brainstem. J. Neurosci. 31, 14037–14045 (2011).

15. Sholl, D. a. Dendritic organization in the neurons of the visual and motor cortices of the cat. J. Anat. 87, 387–406.1 (1953).

16. Ferriera, T. A. et al. Neuronal morphometry directly from bitmap images. Nat. Methods 11, 981–984 (2014).

17. Bonder, D. E. & McCarthy, K. D. Astrocytic Gq-GPCR-Linked IP3R-Dependent Ca2+ Signaling Does Not Mediate Neurovascular Coupling in Mouse Visual Cortex In Vivo. J. Neurosci. 34, 13139–13150 (2014).

18. Martin-Fernandez, M. et al. Synapse-specific astrocyte gating of amygdala-related behavior. Nat. Neurosci. 20, 1540–1548 (2017).

19. Adamsky, A. et al. Astrocytic Activation Generates De Novo Neuronal Potentiation and Memory Enhancement. Cell 174, 59–71 (2018).

20. Maniscalco, J. W. & Rinaman, L. Vagal Interoceptive Modulation of Motivated Behavior. Physiology 33, 151–167 (2018).

21. Roman, C. W., Sloat, S. R. & Palmiter, R. D. A tale of two circuits: CCKNTSneuron stimulation controls appetite and induces opposing motivational states by projections to distinct brain regions. Neuroscience 358, 316–324 (2017).

22. Carter, M. E., Soden, M. E., Zweifel, L. S. & Palmiter, R. D. Genetic identification of a neural circuit that suppresses appetite. Nature 503, 111–4 (2013).

23. Atasoy, D., Betley, J. N., Su, H. H. & Sternson, S. M. Deconstruction of a neural circuit for hunger. Nature 488, 172–177 (2012).

24. Fuente-Martín, E. et al. Leptin Regulates Glucose and Glutamate Transporters in Hypothalamic Astrocytes. J. Clin. Invest. 122, 3900–3913 (2012).

25. Reiner, D. J. et al. Astrocytes Regulate GLP-1 Receptor-Mediated Effects on Energy Balance. J. Neurosci. 36, 3531–3540 (2016).

26. Douglass, J. D., Dorfman, M. D., Fasnacht, R., Shaffer, L. D. & Thaler, J. P. Astrocyte IKKβ/NF-κB signaling is required for diet-induced obesity and hypothalamic inflammation. Mol. Metab. 6, 366–373 (2017).

27. Matott, M. P., Kline, D. D. & Hasser, E. M. Glial EAAT2 regulation of extracellular nTS glutamate critically controls neuronal activity and cardiorespiratory reflexes. J. Physiol. 595, 6045–6063 (2017).

28. Vance, K. M., Rogers, R. C. & Hermann, G. E. PAR1-Activated Astrocytes in the Nucleus of the Solitary Tract Stimulate Adjacent Neurons via NMDA Receptors. J. Neurosci. 35, 776–785 (2015).

29. Treece, B. R., Covasa, M., Ritter, R. C. & Burns, G. A. Delay in meal termination follows blockade of N-methyl-D-aspartate receptors in the dorsal hindbrain. Brain Res. 810, 34–40 (1998).

30. Schindelin, J. et al. Fiji: An open-source platform for biological-image analysis. Nat. Methods 9, 676–682 (2012).

